# Chaperone isoform and interactome mapping reveals functional diversification of DNAJA2–DNAJA4 complexes via stress-regulated isoforms

**DOI:** 10.64898/2026.09.23.753896

**Authors:** Tahani Kadah, Nadeen Akaree, Anastasya Brodov-Nevo, Anatoly Meller, Flonia Levy-Adam, Reut Shalgi

## Abstract

The human HSP70 chaperone network maintains cellular proteostasis through a diverse repertoire of HSP70s and co-chaperones. Here we examine alternative isoforms and co-chaperone hetero-complexes as additional sources of network complexity. To that end, we systematically mapped the isoform, tissue-expression, and interaction landscapes of the human HSP70 network, revealing a modular interactome containing known and novel DNAJ-DNAJ interactions. We found both tissue-expression isoform divergence as well as widespread alternative-canonical isoform co-expression, suggesting additional modes of functional diversification. Focusing on the uncharacterized DNAJA2–DNAJA4 hetero-complex, we identified the stress-inducible isoform DNAJA4-CTD-II. DNAJA4-CTD-II formed hetero-complexes with DNAJA2 and DNAJA4, with both interactions enhanced following sodium arsenite stress. Functionally, DNAJA4-CTD-II co-localized with TDP-43 aggregates and significantly suppressed their accumulation in a DNAJA2-dependent manner. Together, our data reveal extensive, uncharted isoform and interaction complexity within the HSP70 network, and uncover isoform-dependent hetero-complex remodeling as a new layer of chaperone network regulation.

## Introduction

The molecular chaperone network is a central component of the proteostasis system ^1^. Chaperones facilitate proper protein folding by recognizing misfolded or non-native polypeptides, unfolding them when necessary, and enabling their refolding^1^. HSP70 chaperones, the most ubiquitous chaperone family^2^, require the association of two additional class members of co-chaperones to exert their chaperoning function: DNAJ proteins (HSP40) and nucleotide exchange factors (NEFs) ^3^. DNAJs promote HSP70’s ATPase activity, and thereby it’s client binding, whereas NEFs promote ADP exchange and release of unfolded clients ^3^. While the HSP70 cycle is highly conserved in evolution, the chaperone families have undergone major expansions ^4–6^. The human genome encodes 13 HSP70s, 13 NEFs, and nearly 50 DNAJ proteins ^5^. The diversity of human DNAJ proteins has been proposed to regulate multiple aspects of HSP70 functions: client specificity, subcellular localization, and more ^3–5^, leading to functional diversification of the HSP70 chaperone network.

DNAJ proteins can be classified into three classes ^7^, while class A and class B DNAJs are known to form homodimers ^4,5^. Several studies demonstrated heterodimerization between specific DNAJ proteins, including DNAJA2–DNAJB1 and DNAJB12–DNAJB14, while interactome mapping studies suggested additional DNAJ-DNAJ interactions ^8–11^. These observations indicate that DNAJ proteins can assemble into selective hetero-complexes with potentially distinct functions. However, the prevalence and functional significance of such complexes remain largely unknown.

Alternative isoforms provide an additional source of DNAJ diversification. Previous studies showed that isoforms of DNAJB12, DNAJB14, DNAJB6 and DNAJB2 differ in localization and aggregation-modulating activity ^10,12–14^. However, the landscape of DNAJ isoforms and their contribution to chaperone function remain unexplored.

Together, hetero-complex formation and alternative isoforms have the potential to considerably expand the combinatorial organization of the HSP70 network, with potentially diverse functions in proteostasis regulation. Yet these two sources of diversity have largely been studied independently, with only a handful of individual examples characterized, and the mutual effects of one on the other has not been examined. Thus, their combined contribution to chaperone network organization remains unknown.

Here we performed a systematic interactome characterization of the HSP70 subnetwork, using the LUMIER assay for pairwise chaperone interactions. This interaction network mapping revealed broad, yet selective, chaperone modules and hetero-complexes that include multiple DNAJs. Expanding on chaperone isoforms, we additionally identified dozens of chaperone isoforms with interesting tissue expression patterns. Focusing on the previously uncharacterized hetero-complex of DNAJA4 and DNAJA2, we found that a short isoform of DNAJA4, termed DNAJA4-CTD-II, was highly induced in response to specific stress conditions. DNAJA4-CTD-II was able to form hetero-complexes with the full length DNAJA2 and DNAJA4, and complex formation increased in stress. Finally, we found that DNAJA4-CTD-II highly and specifically localized to TDP-43 aggregates and was able to suppress TDP-43 aggregation, in a DNAJA2-dependent manner. Our data provides a comprehensive map of hetero-complexes and isoforms for the HSP70 network, establishing that isoform-involving hetero-complex formation expands the functional versatility of the chaperone network.

## Results

### Chaperone isoforms show diverse expression patterns across human tissues

The human HSP70 chaperone network has undergone substantial evolutionary expansion, with 13 HSP70 proteins, 13 NEFs, and 49 members of the DNAJ family. Beyond individual family members, this diversity can potentially be further amplified at the transcript level, through isoform diversity. To systematically assess this layer of complexity, we mapped naturally occurring mRNA isoforms across these chaperone families by integrating several transcript annotation and domain mapping databases (see Methods, Table S1). Our analysis revealed a distribution of isoform numbers for all three families, with 60%, 88% and 85% of HSP70s, DNAJs and NEFs (respectively) having more than one isoform (Fig. 1A).

**Figure 1:**
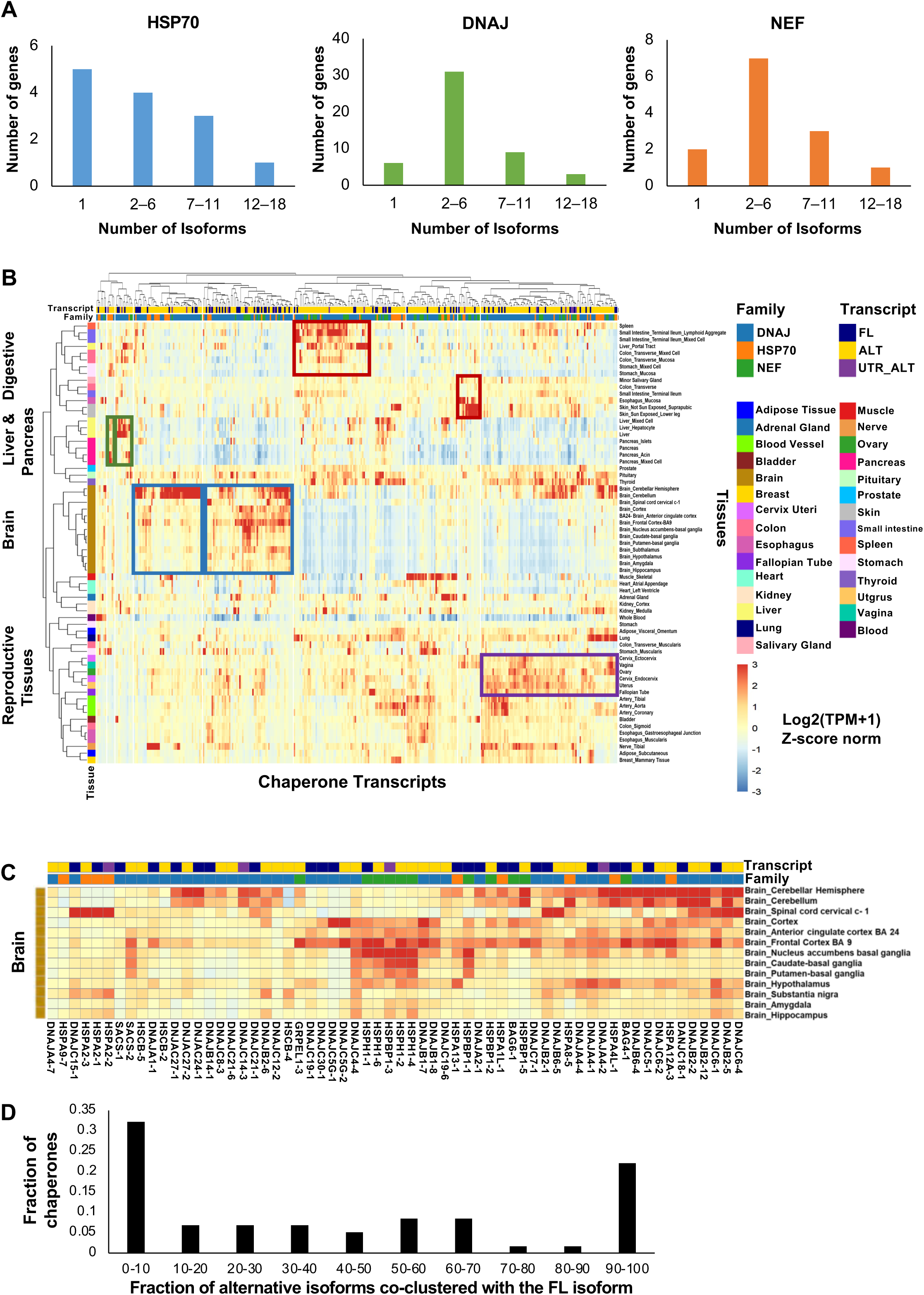
Diverse chaperone isoform expression patterns across human tissues. (A) Distribution of isoform numbers per gene across the HSP70, DNAJ, and NEF chaperone families. Naturally occurring mRNA isoforms were systematically mapped by integrating GENCODE transcript annotations, accessed through UCSC and Ensembl, with protein domain information from UniProt, Pfam, and InterPro (see Methods). (B) Heatmap of transcript expression levels across human tissues (GTEx), shown as log_2_(TPM+1), z-score normalized values for isoforms of the HSP70, DNAJ, and NEF families. Transcripts are color coded by chaperone family and isoform type, either full length (FL) or alternative, which can have a different coding sequence relative to the FL (ALT), or only different UTR relative to the FL (UTR_ALT). Hierarchical clustering was performed on transcripts and tissues to reveal patterns of expression and clusters across tissues. (C) Close up on the Brain-wide expression cluster. (D) Co-expression analysis of all alternative isoforms with their corresponding full-length (FL) isoform. For each chaperone, the fraction of alternative isoforms co-clustering with its FL isoform within the same expression cluster was calculated.

Next, we analyzed isoform-level expression across human tissues using the GTEx transcriptomics database ^15^. Isoforms were classified into full-length (FL) and alternative (ALT) according to their domain composition (see Methods, Table S1). Clustering analysis revealed extensive variation in isoform expression across tissues (Fig. 1B). Tissue grouping according to expression profiles displayed distinct clusters, many corresponding to major tissue types. Two distinct, yet adjacent, brain/CNS specific clusters emerged, one showing preferentially higher chaperone expression throughout brain regions and the other more specific to the cerebellum (Fig. 1B, blue). In addition, the analysis identified two digestive system clusters (Fig. 1B, red), and two adjacent liver-pancreas clusters (Fig. 1B, green). Reproductive tissues also clustered together (Fig. 1B, purple), albeit chaperones in this cluster had a broader expression pattern across many tissues.

Interestingly, the brain-wide preferential cluster exhibited prominent expression of multiple canonical full-length chaperone isoforms, alongside some of their alternative isoforms (Fig. 1C). Those included the three DNAJ class A members DNAJA1, DNAJA2, DNAJA4, the neuronal chaperone DNAJB2 ^16^, DNAJB14, which was previously shown to regulate ALS-associated FUS aggregation ^10^, and the chaperone DNAJC7, mutations in which were found in some cases of ALS ^17^. In addition, this cluster contained the full-length isoforms of the HSP70s HSPA2 and HSPA13, and the NEFs HSPH1 ^18^ and BAG6. In contrast, the cerebellum specific cluster mainly included non-canonical alternative chaperone isoforms.

We then asked if alternative chaperone isoforms tended to undergo functional diversification in their tissue expression patterns. We examined the co-expression of different isoforms of the same chaperone, which often segregated into distinct expression clusters (Fig. 1B). We tested whether alternative isoforms and their corresponding FL isoform co-clustered together in tissue expression. For each chaperone, we calculated the fraction of alternative isoforms that were assigned to the same expression cluster as the canonical FL isoform. The resulting U-shaped distribution revealed a substantial proportion of chaperones showing either complete divergence, i.e. none of the alternative isoforms co-clustered with the FL, or full concordance, i.e. all alternative isoforms co-clustered with the FL (Fig. 1D, Table S2). Among the 59 chaperones with multiple isoforms, 58% exhibited substantial divergence in tissue-expression patterns, with more than half of their alternative isoforms assigned to different expression clusters than the canonical FL isoform (Fig. 1D). Nevertheless, the remaining 42% retained co-expression with most of their alternative isoforms. Therefore, while tissue-specific expression diversification appears to be a major source of isoform functional diversification, the widespread co-expression of many alternative isoforms with their canonical counterparts suggested that additional mechanisms contribute to isoform specialization within the same cells.

### Mapping chaperone interactome modules and hetero-complexes through LUMIER profiling

We next sought to systematically define the core interaction architecture underlying the HSP70 chaperone network. To that end, we mapped pairwise interactions among the canonical FL isoforms within the HSP70 network, with a few chaperones from other families (see Methods), using the LUMIER assay ^19^. In this assay, FLAG-tagged bait chaperones were co-expressed together with each of a panel of Nanoluciferase-tagged prey chaperones in HEK293T cells, while interaction strength was quantified by measuring luciferase activity associated with the immunoprecipitated bait, and normalized to bait levels (see Methods). The resulting interaction scores were further z-score normalized along the prey axis to highlight the strongest preferential binders for each prey chaperone. Accordingly, the map is not expected to be symmetric, although for many cases symmetry is indeed observed. The resulting interactome map revealed a structured, fairly sparse, network of associations among DNAJ proteins, HSP70s, and NEFs (Fig. 2A, Table S3). Unsupervised clustering identified discrete interaction modules, which were in some cases interconnected, indicating that the chaperone network is organized into specific preferential interaction subnetworks rather than forming a uniformly connected system.

**Figure 2:**
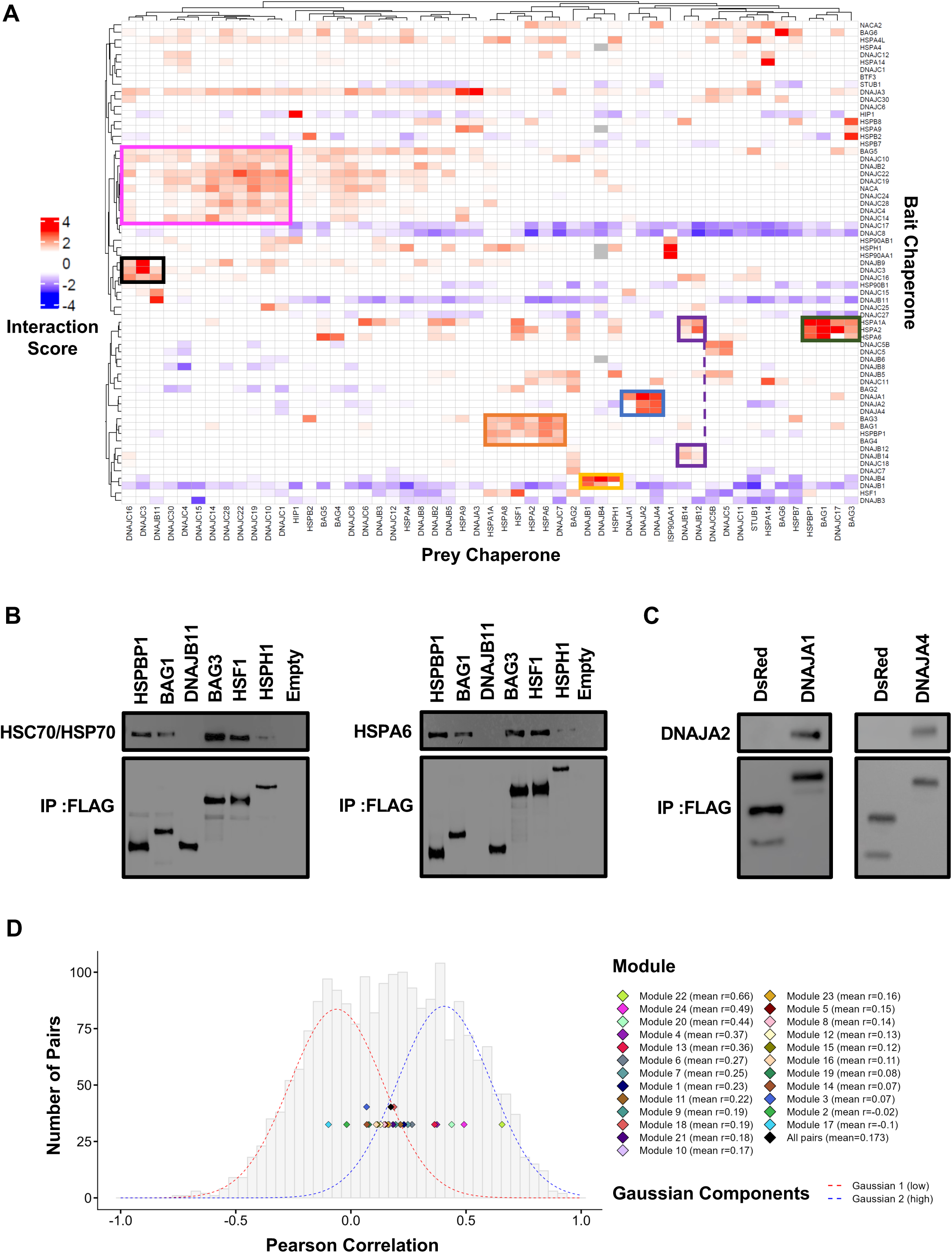
Mapping interaction modules and hetero-complex architecture within the HSP70 chaperone network. (A) Heatmap of protein-protein interactions measured using the LUMIER assay. FLAG-tagged bait chaperones were co-expressed with NanoLuciferase-tagged prey chaperones, followed by immunoprecipitation and luminescence-based detection. Interaction scores were then calculated by normalizing luminescence to bait levels, measured using FLAG ELISA (see Methods). Colors represent interaction scores (z-score normalized per prey), scores between -1 and 1 were set to white. Hierarchical clustering of bait and prey chaperones revealed patterns of interaction across the chaperone network. Selected interaction modules are marked by colored frames. (B) Validation of selected interactions identified in the LUMIER assay, using co-immunoprecipitation (co-IP). FLAG-tagged proteins were immunoprecipitated, and endogenous interacting proteins were detected by immunoblotting using anti-HSPA6 (right) or anti-HSC70/HSP70 (left) antibodies. Representative blot out of 2 independent biological replicate experiments. Quantification shown in Fig. S1A. (C) as in (B) using anti-DNAJA2 antibody. Representative blot out of 2 independent biological replicate experiments. Quantification shown in Fig. S1B. (D) Distribution of Pearson correlation coefficients calculated from expression data (GTEx) for all transcripts corresponding to the canonical isoforms of the chaperones included in the LUMIER interaction map. The background histogram (grey) represents the distribution of all pairwise correlations. A Gaussian mixture model (GMM) was fitted to the distribution, revealing a bimodal distribution: a low-correlation population (red dashed gaussian) and a high-correlation population (blue dashed gaussian). Colored diamonds denote the mean pairwise correlation values for individual LUMIER-defined modules (see Table S3). A small subset of modules shows co-expression correlation values within the high component, consistent with coordinated expression across tissues. Notably, Module 22 (DNAJB12/DNAJB14) and Module 24 (DNAJA2/DNAJA4) exhibit the highest mean correlations.

For most DNAJs, homodimerization was indeed detected in our assay (in 22 out of 31 DNAJs examined). The clustered map additionally highlighted several hetero-complexes between different DNAJs. For example, DNAJA1, DNAJA2, and DNAJA4 were found to preferentially co-interact (Fig. 2A, blue module), consistent with previous indications from a proteomics-based dataset ^11^. A preferential binding was found between DNAJB14 and DNAJB12, which additionally interacted with the three cytosolic HSP70s HSPA1A, HSPA2 and HSPA6, and more weakly with their homolog DNAJC18 (purple module). We previously characterized the DNAJB12-DNAJB14-HSP70 hetero-complex as a strong aggregation suppressor of the ALS-associated mutant FUS ^10^, assigning a specific function to that module. Interestingly, some of the modules involved specific DNAJ-NEF interactions, revealing binding preferences across co-chaperone families. For example, DNAJB4 and DNAJB1 strongly co-interacted, consistent with previous proteomics-based studies ^11^, and demonstrated preferential binding to the NEF HSPH1 (Fig. 2A, yellow module). Previous studies found that HSPH1 cooperated with DNAJB1 and HSP70 as part of the mammalian disaggregation machinery, demonstrating a specific function for this interaction ^20^. In another module, DNAJC7, which co-clustered with several cytosolic HSP70s, robustly interacted with the NEFs BAG1 and BAG3, in agreement with previous BAG3 IP-MS studies ^21^, in addition to BAG4 and HSPBP1 (Fig. 2A, orange module). In addition, our analysis uncovered previously uncharacterized preferential associations, such as the module consisting of preferential interactions amongst the ER resident chaperones DNAJC3, DNAJC16, DNAJB9 and DNAJB11 (black module). Beyond the well-defined small modules, the network exhibited one broad module enriched for many C class DNAJ proteins, probably representing a sub-network of interactions (Fig. 2A, pink box).

To validate the interaction patterns identified in our LUMIER interactome mapping, we performed co-immunoprecipitation (co-IP) experiments for selected protein pairs (Fig. 2B,C, S1). The interactions between DNAJB4 and HSPH1, as well as between DNAJB4 and DNAJB1, were validated using GFP- and FLAG-tagged constructs (Fig. S1C,D). We confirmed multiple interactions identified in the screen between NEFs and the endogenous HSP70 family members HSPA8 and HSPA1A (aka HSC70/HSP70), as well as HSPA6, while DNAJB11 was confirmed as non-interactor (Fig. 2B, S1A). In addition, we also verified the interactions between DNAJA4 or DNAJA1 and the endogenous DNAJA2 (Fig. 2C, S1B).

The LUMIER interactome mapping revealed many modules and hetero-complexes, demonstrating another aspect of inter-family functional diversification. Yet since it was performed using exogenously expressed proteins, the identified modules represent potential hetero-complexes, which are contingent on the chaperones in the module/hetero-complex being co-expressed within the same tissue or cell type. To further investigate whether chaperones within interaction modules are coordinated in their tissue expression profiles, we again turned to the GTEx human tissue expression database and examined the correlations between proteins identified by the interactome map as belonging to the same modules. Pearson correlations were calculated for each module/hetero-complex (see Methods) and compared to the distribution of pairwise correlations between all chaperones in the interactome map. The underlying distribution of pairwise correlation coefficients was bimodal, with one mode centered around 0.4 and the other around zero (Fig. 2D). Out of 24 modules/hetero-complexes examined, most fell between the two gaussian components, suggesting that members of a given module are co-expressed in overlapping, but not identical, subsets of tissues. Interestingly, only five modules exhibited increased co-expression across tissues, belonging to the second component (Fig. 2D). The highest co-expression correlation was observed for module 22, comprising DNAJB14 and DNAJB12, for which we previously assigned a function in FUS aggregation suppression ^10^. The second highest module was hetero-complex 24, consisting of DNAJA4 and DNAJA2, both of which showed preferential expression mostly in the brain, but its functional impact was not examined before. We therefore next sought to investigate the potential function and regulation of this hetero-complex.

### Isoform specific characterization of DNAJA2 and DNAJA4 reveals novel hetero-complexes

The coordinated expression and interaction profile of DNAJA2 and DNAJA4 prompted us to investigate this module in greater depth. We additionally aimed to determine how isoform diversification within this module can potentially fine-tune the function of this hetero-complex.

Based on our mapping above, we identified multiple naturally occurring variants of DNAJA2 and DNAJA4. DNAJA4 encodes 12 isoforms, while DNAJA2 encodes 4 isoforms. We focused on isoforms with distinct domain composition, in order to investigate main structure-function relationships in naturally occurring isoforms. Accordingly, for DNAJA4 we selected the full-length DNAJA4 (-FL), an alternative transcription start isoform which contains the CTD-II, a domain known to be involved in substrate binding ^5^, and the dimerization domain (DD), and an isoform that contains the J-Domain with GF linker (Fig. 3A). For DNAJA2 we selected isoforms representing the canonical full-length protein, and a C-terminal truncated isoform which additionally had an alternative promoter, that contains the CTD-I (Fig. 3A).

**Figure 3:**
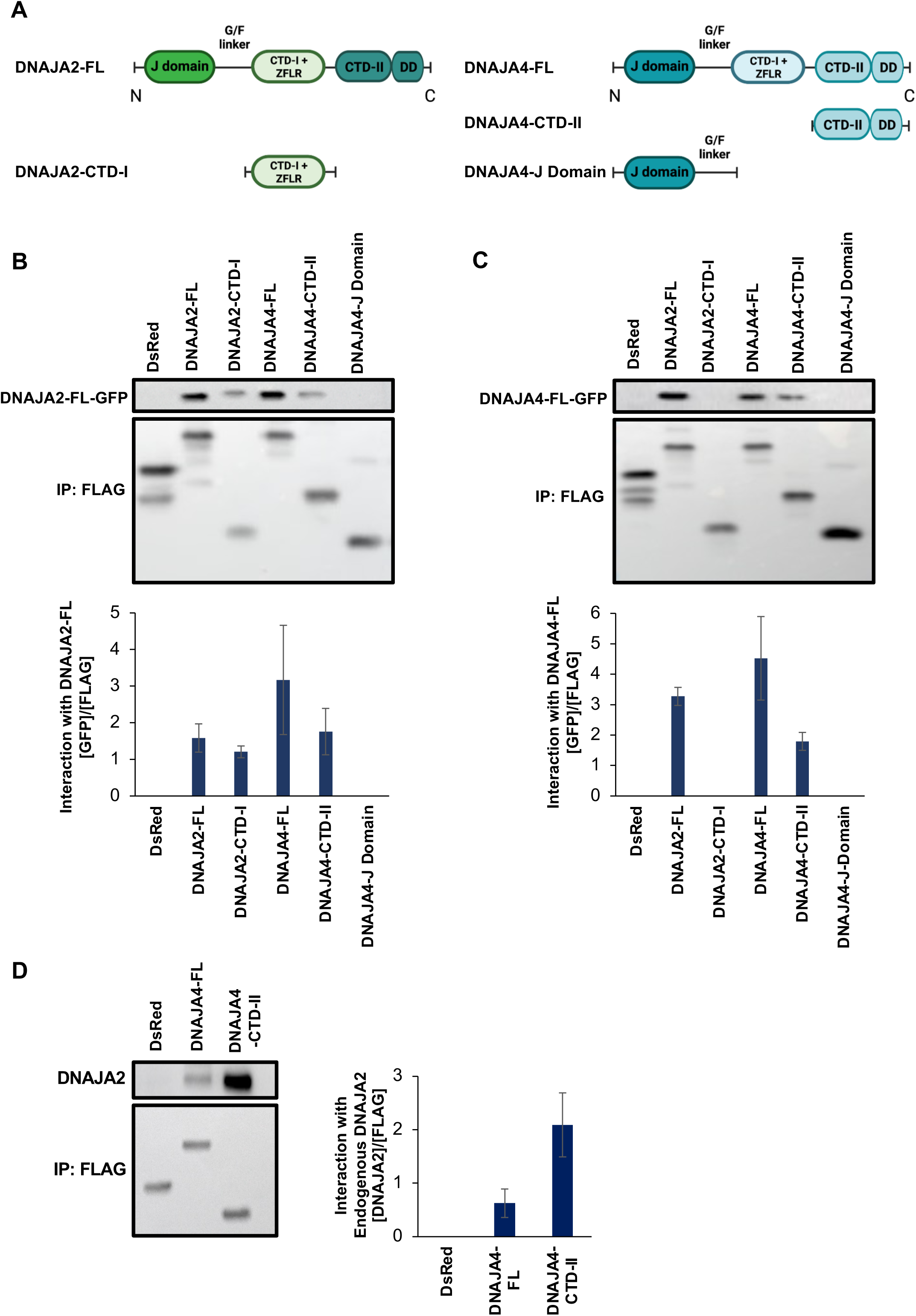
DNAJA4 and DNAJA2 form hetero-complexes with their naturally occurring isoforms. (A) Schematics of DNAJA2 and DNAJA4 protein isoforms examined. DNAJA4-FL: ENST00000343789.7, DNAJA4-CTD-II: ENST00000493321.1, DNAJA4-J-Domain: ENST00000423642, DNAJA2-FL: ENST00000317089.10, DNAJA2-CTD-I: ENST00000617000.1. (B) DNAJA2-FL-GFP and (C) DNAJA4-FL-GFP interactions with different FLAG tagged isoforms of DNAJA2 and DNAJA4 proteins were assayed using co-IP. Bar plots represent the relative interaction, calculated by dividing the quantified GFP signal by the quantified FLAG signal of the pulldown protein. Representative blot is shown for N=3 independent biological replicates, plots presented as mean+/-SEM. Inputs presented in Fig. S2A,B for panels (B) and (C), respectively. (D) Endogenous DNAJA2 interacts with FLAG tagged DNAJA4-FL and DNAJA4-CTD-II isoforms. Representative blot is shown for N=2 independent biological replicates, plots represent mean +/- STD. Input is presented in Fig. S2C.

We performed Co-IP experiments to examine the full-length proteins and to assess potential interactions of their isoforms. FL proteins were tagged with GFP, while other isoforms were tagged with FLAG, and pulldown was performed using anti-FLAG beads. Our results showed that DNAJA2-FL formed homodimers, as expected, and interacted with DNAJA2-CTD-I to form a hetero-complex (Fig. 3B, S2A). DNAJA2-FL also associated with DNAJA4-FL, forming a hetero-complex, as found in the LUMIER interactome above, and with the DNAJA4 isoform DNAJA4–CTD-II (Fig. 3B, S2A). Similarly, DNAJA4-FL formed homodimers and its reciprocal interaction with DNAJA2-FL was further confirmed (Fig. 3C, S2B). DNAJA4-FL also interacted with DNAJA4–CTD-II, but not with the DNAJA4-J-domain isoform or DNAJA2-CTD-I (Fig. 3C, S2C). While DNAJA4 is very lowly expressed in the cells used (HEK293T cells), DNAJA2 is well expressed, and we thus continued to examine the endogenous DNAJA2 and its potential interactions with FLAG-tagged DNAJA4 isoforms. Co-IP showed that both the FL and the CTD-II DNAJA4 isoforms indeed interacted with the endogenous DNAJA2 (Fig. 3D, S2C).

### The novel isoform DNAJA4-CTD-II is highly induced in response to heat shock and sodium arsenite stresses

The expression patterns of these alternative isoforms across tissues partially overlapped with that of the full-length isoforms, while DNAJA4-J-Domain was nearly absent throughout (Fig. S3A). We next asked whether these isoforms might be inducible by stress. To address this, we designed isoform specific qPCR primers for each of the isoforms examined, with the exception of the DNAJA4-J Domain, for which isoform specific primers could not be designed. We then performed qPCR in response to different proteotoxic stress conditions. To examine potential stress inducibility of DNAJA4, we used MCF-7 cells, as these cells express DNAJA4-FL basally, whereas other cell lines show much lower levels of DNAJA4 expression (Fig. S3B). Notably, using genomic DNA as a standard, we were able to perform quasi-absolute qPCR, allowing for comparison between isoforms of the same gene (see Methods). MCF-7 cells were subjected to a variety of stress conditions, including ER stress (Thapsigargin, TG, 1uM for 2h), oxidative stress (sodium arsenite, 200uM, for 2h), proteasome inhibition (MG-132, 1uM, for 4h), and heat shock (44^0^C, 2h). Both DNAJA2-FL and DNAJA2-CTD-I exhibited relatively stable expression in response to the different stress conditions (Fig. 4A,B). While DNAJA2-CTD-I showed a somewhat stronger induction than DNAJA2-FL under proteasome inhibition, for other stresses, the overall ratio between DNAJA2-CTD-I and DNAJA2-FL remained largely unchanged (Fig. 4B).

**Figure 4:**
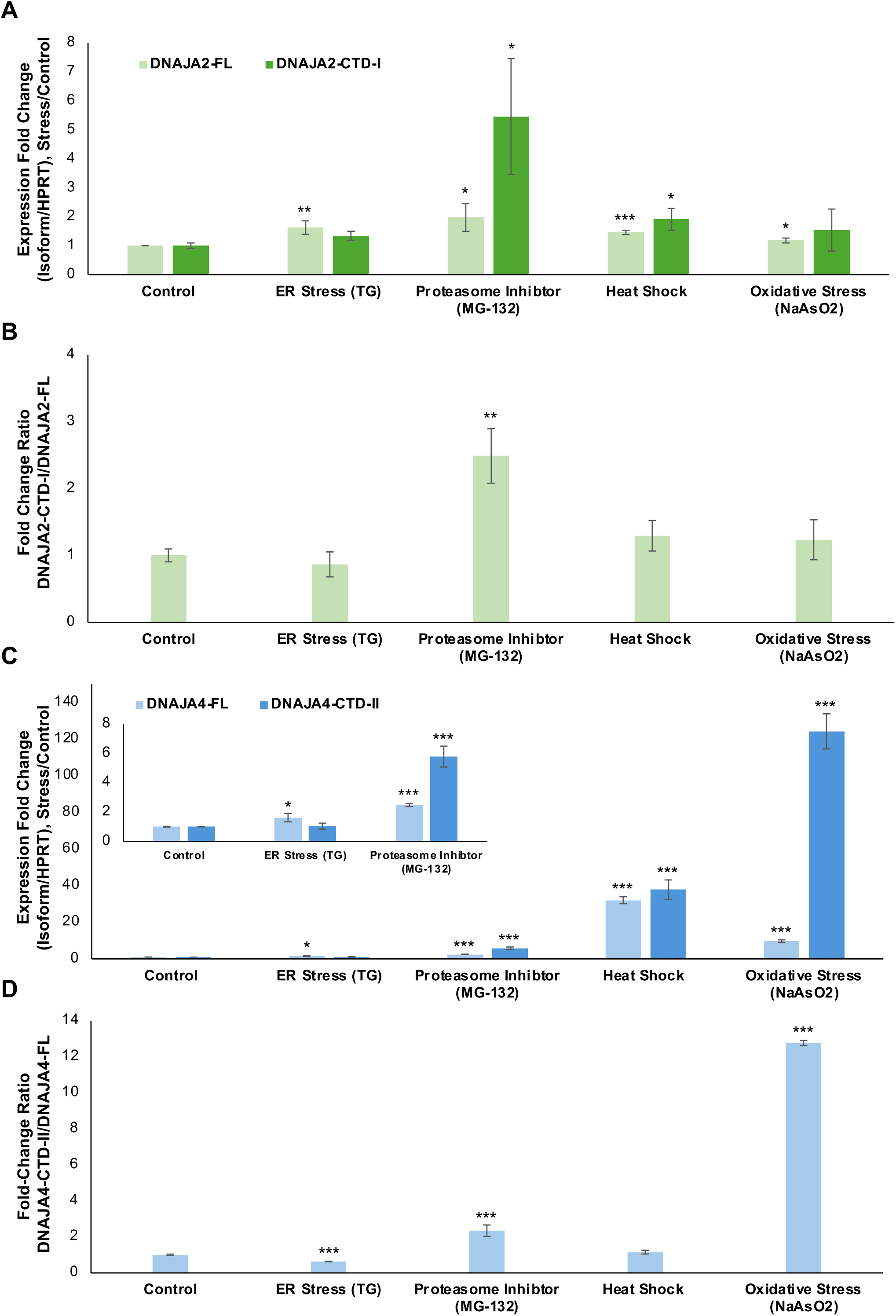
DNAJA4-CTD-II expression is highly induced in response to stress. (A) Expression of DNAJA2 isoform across different stress conditions (TG, 1uM for 2h; MG-132, 1uM for 4h; heat shock, 44°c for 2h; sodium arsenite, 200uM for 2h). Isoforms expression is shown as the ratio of isoform copy number (determined using a genomic DNA standard, see Methods) to HPRT copy number, and further normalized to the non-stressed condition. Data shown as mean ± SEM of independent biological replicates (Non-stressed, N=6; TG, N=3; MG-132, N=4; sodium arsenite, N=3; heat shock, N=6). (B) Fold change ratio of DNAJA2-CTD-I relative to the full-length (FL) isoform, corresponding to data shown in (A). (C) As in (A), for DNAJA4 isoforms. The inset shows zoom-in of ER stress and proteasome inhibitor stress. Data presented as mean ±SEM from independent biological replicates, as in panel (B). (D) As in (B) for DNAJA4 isoforms. Statistical significance: *p < 0.05, **p < 0.01, ***p < 0.001, calculated using two-tailed t-test.

In contrast, both DNAJA4-FL and DNAJA4-CTD-II were markedly induced in response to stress (Fig. 4C). In response to MG-132 treatment DNAJA4-FL was modestly induced (∼2.5 fold), while DNAJA4-CTD-II was upregulated about 6 fold. Notably, both DNAJA4-FL and DNAJA4-CTD-II were vastly induced in response to heat shock and sodium arsenite stresses (Fig. 4C). Strikingly, DNAJA4-CTD-II showed a substantially stronger upregulation than DNAJA4-FL particularly under sodium arsenite treatment, as can be appreciated from the DNAJA4-CTD-II/DNAJA4-FL expression ratio (Fig. 4D). Although both isoforms were highly upregulated by stress, the proportional rise in DNAJA4-CTD-II expression in response to sodium arsenite suggested that different stresses may differentially modulate isoform usage.

### DNAJA4-CTD-II hetero-complexes interactions are enhanced during sodium arsenite stress

Due to the strong upregulation of DNAJA4 isoforms in sodium arsenite stress we asked whether this condition additionally influences the interactions among the different DNAJA isoforms. We performed co-IP in cells co-expressing FLAG-tagged isoforms of DNAJA4 or DNAJA2 together with GFP-tagged DNAJA4-FL (Fig. 5A) or DNAJA2-FL (Fig. 5B), before or after sodium arsenite treatment for 2h followed by 2h recovery. While under normal conditions DNAJA4-FL formed homodimers and interacted with DNAJA2 and with DNAJA4-CTD-II (Fig. 5A) as shown above (Fig. 3B,C), upon arsenite exposure, we observed a distinct, isoform-specific effect, whereby DNAJA4-CTD-II showed a markedly increased interaction with DNAJA4-FL (Fig. 5A, S4A). We note that these interactions were measured for exogenous proteins and are therefore independent of any potential induction in endogenous DNAJA4-CTD-II shown above. The homo-interactions of DNAJA4-FL with itself, as well as its interaction with DNAJA2-FL remained largely unchanged following arsenite stress (Fig. 5A,B). Interestingly, DNAJA2-FL also showed a pronounced increase in its interaction with DNAJA4-CTD-II during arsenite treatment (Fig. 5B, S4B), suggesting that arsenite promotes DNAJA4-CTD-II hetero-complex formation.

**Figure 5:**
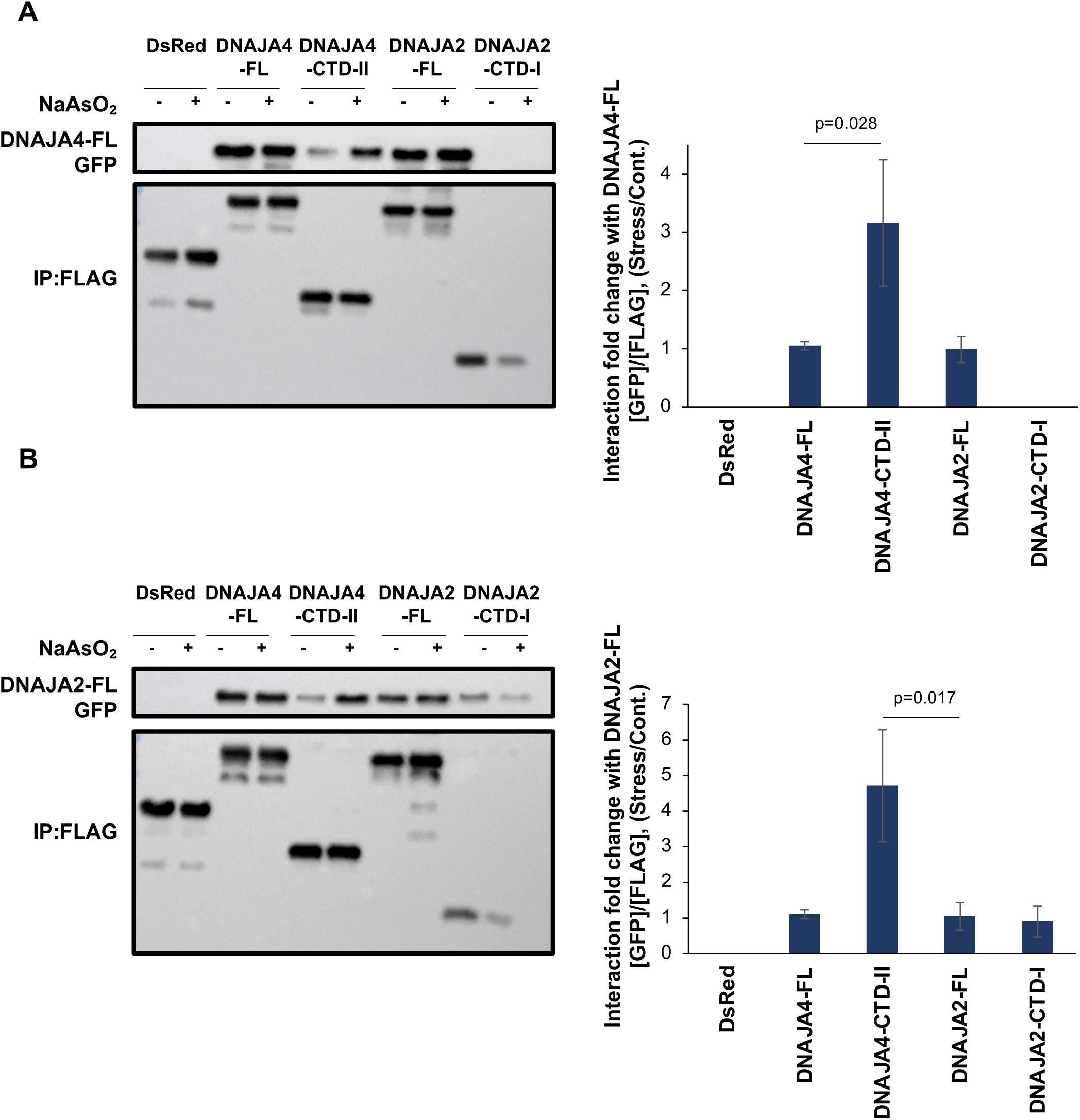
Arsenite stress enhanced interactions between DNAJA4-CTD-II and the full-length DNAJA4 and DNAJA2. (A-B) Co-IP showing interactions between DNAJA4-FL–GFP (A) and DNAJA2-FL-GFP (B) with FLAG-tagged isoforms of DNAJA4 and DNAJA2, in arsenite stress. HEK293T cells co-transfected with GFP-tagged full length DNAJA4/DNAJA2 and FLAG-tagged DNAJA2 or DNAJA4 isoforms, were treated with sodium arsenite (200 µM) for 2 h followed by a 2 h recovery, and then subjected to co-IP using anti-FLAG antibody coated beads. Interactions were detected using an anti-GFP antibody, and anti-FLAG immunoblotting was used to normalize for the amount of immunoprecipitated isoform. Band intensities were quantified using Fiji. Interaction fold changes were calculated by normalizing GFP signal intensity to FLAG signal intensity under stressed conditions and dividing this ratio by the one in the corresponding non-stressed condition. Data presented as mean ± SEM from N = 3 independent biological replicates. DNAJA4-CTD-II showed a marked increase interaction with DNAJA4-FL (A) as well as DNAJA2-FL (B) upon sodium arsenite treatment. Inputs presented in Fig. S4. Statistical significance calculated using two-tailed t-test.

### DNAJA4 isoforms modulate TDP-43 aggregation in a DNAJA2-dependent manner

Having characterized the interaction profiles of the different DNAJA4 and DNAJA2 isoforms, we next aimed to determine how these different isoforms, and their hetero-complexes, contribute to proteostasis regulation. We sought to examine whether the individual isoforms of DNAJA2 and DNAJA4 may contribute to protein aggregation regulation, specifically, the aggregation of the ALS-causative protein TDP-43. We used the aggregation-prone variant TDP-43–ΔNLS-2KQ-YFP ^22^, which had a mutated nuclear localization signal (NLS) and carries acetylation-mimic mutations. Indeed, while WT-TDP-43 was nuclear, TDP-43-ΔNLS–2KQ was largely cytosolic, and tended to form aggregates (Fig. 6A, S5A). To test the ability of each DNAJA2/DNAJA4 isoform to modulate the aggregation of TDP-43, we quantified the extent of TDP-43-ΔNLS–2KQ aggregation using the FACS-based PulSA method ^23^ (Fig. S5B), as we previously performed for FUS aggregation ^10^. We used the aggregation modulation score, defined as the log2 fold change in the percentage of aggregate-containing cells (AGG+) in the presence of each modulator compared to the respective control (co-expression of an unrelated protein, DsRed), and significance was assessed using the 95% confidence interval (see Methods). The positive control DNAJB5, previously shown to suppress TDP-43-ΔNLS–2KQ aggregation ^24^, indeed gave a significant aggregation modulation score, whereas co-expression of the chaperone DNAJA1 showed no effect on TDP-43 aggregation (Fig. 6B, S5B). DNJA2-FL also significantly suppressed TDP-43-ΔNLS–2KQ aggregation, consistent with recent reports ^25,26^, and DNAJA2-CTD-I had a substantially reduced ability to suppress aggregation (Fig. 6B, S5B). Interestingly, we found that DNAJA4 isoforms also significantly reduced TDP-43 aggregation, while DNAJA4-CTD-II showed a stronger effect compared to DNAJA4-FL (Fig. 6B).

**Figure 6:**
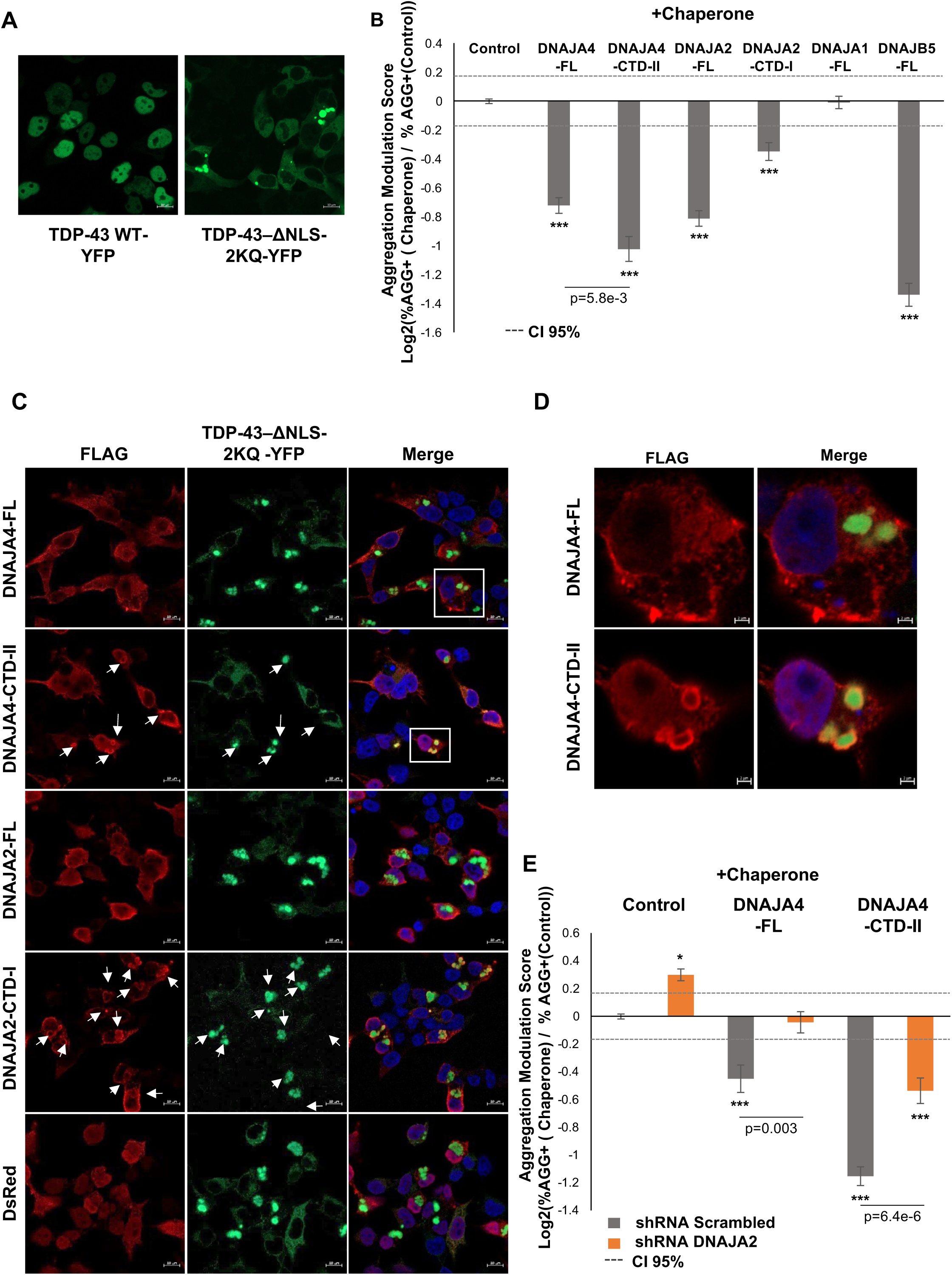
Modulation of TDP-43 aggregation by DNAJA4 and DNAJA2 isoforms. (A) Fluorescence microscopy images of HEK293T cells transfected with TDP-43-WT-YFP or TDP-43–ΔNLS– 2KQ-YFP, 48h post transfection, scale bar, 10 μm, see also Fig. S5A (B) Aggregation modulation scores of TDP-43, in HEK293T cells co-expressing TDP-43–ΔNLS–2KQ-YFP together with isoforms of DNAJA2 or DNAJA4 (or DsRed as control). The fraction of cells containing TDP-43 aggregates (AGG+) was quantified using PulSA and normalized to the corresponding DsRed co-expression control. Values represent log2 fold changes in the percentage of AGG+ cells relative to the control, as previously described ^10^, see Methods. Multiple DsRed controls (n = 28) were used to calculate the 95% confidence interval (CI 95%, dashed grey line), defined as 2*STD of the control population. Negative values indicate a reduction in the proportion of AGG+ cells, reflecting suppression of TDP-43 aggregation. Data presented as mean ± SEM of N=7/7/7/7/5/5/3 independent biological experiments for DsRed/DNAJA4-FL/DNAJA4-CTD-II/DNAJA2-FL/DNAJA2-CTD-I/DNAJA1-FL/DNAJB5-FL, respectively, with total replicates of n=30/19/20/17/14/14/12/9. *-p<0.05, ***-p<0.003, empirical p-values, see Methods. DNAJA4-CTD-II exhibited a significantly stronger suppression of TDP-43 aggregation compared to DNAJA4-FL (p=5.8e-3), calculated using two-tailed t-test. (C) Confocal immunofluorescence images of FLAG-tagged DNAJA4 isoforms (red), stained using anti-FLAG antibody with a far-red secondary antibody, co-expressed with TDP-43–ΔNLS–2KQ-YFP (green). Nuclei were stained with DAPI (blue). White arrows indicate TDP-43 aggregates co-localized with chaperones. Representative images are shown from N = 2 independent biological experiments, scale bar - 10 μm. Additional images are provided in Fig. S5C. (D) The boxed region from panel (C) are shown in higher magnification, scale bar – 2 μm. (E) Aggregation modulation by DNAJA4 isoforms was calculated as (B) in shRNA Scrambled and shRNA DNAJA2 cell lines. Values were normalized to control + shRNA Scrambled. Data presented as mean ± SEM from N=5/3/4 independent biological experiments for DsRed/DNAJA4-FL/DNAJA4-CTD-II, respectively, with total replicates of n=24/14/18 in shRNA DNAJA2 and n=22/13/17 in shRNA Scrambled. Dashed lines – CI 95%. *-p<0.05, ***-p<0.003, empirical p-values, see Methods. Differences between each pair of shScrambled and shDNAJA2 scores were all significant (p= 1.25e-7, 3.01e-3, 6.4e-6 for Control, DNAJA4-FL and DNAJA4-CTD-II, respectively), calculated using two tailed t-test.

To further explore the potential relationship between these isoforms and TDP-43–ΔNLS–2KQ aggregates, we examined the spatial distribution of TDP-43-ΔNLS–2KQ in cells co-expressing the different DNAJA isoforms using immunofluorescence microscopy. Our experiments revealed distinct localization patterns among the DNAJA2 and DNAJA4 isoforms: while the DNAJA2/A4 full-length isoforms showed a cytosolic diffused pattern with no aggregation localization, DNAJA4-CTD-II and DNAJA2-CTD-I showed clear co-localization with TDP-43 aggregates (Fig. 6C,D, S5C,D). These observations suggest that both the CTD-II and CTD-I regions, previously implicated in substrate recognition ^5^, can specifically associate with aggregated TDP-43, however, DNAJA4-CTD-II had a substantially greater functional effect on TDP-43 aggregation.

Lastly, we asked whether the hetero-complex formation we detected above contributed to the functional effects on TDP-43 aggregation. We further examined the role of the endogenous DNAJA2 in the modulation of TDP-43 aggregation by DNAJA4 isoforms. We generated stable HEK293T cell lines expressing an shRNA to knock down DNAJA2, or a scrambled shRNA as control, with over 40% reduction in endogenous DNAJA2 protein levels (Fig. S5E). We then co-expressed TDP-43-ΔNLS–2KQ and DNAJA4 isoforms in these two cell lines, and assayed aggregation modulation. Our results revealed that endogenous DNAJA2 played a critical role in supporting the function of both DNAJA4-FL and DNAJA4-CTD-II with respect to TDP-43 aggregation suppression (Fig. 6E). Upon knockdown of DNAJA2, TDP-43-ΔNLS–2KQ aggregation mildly but significantly increased (Fig. 6E). Surprisingly, DNAJA4-FL completely lost its ability to suppress TDP-43-ΔNLS–2KQ aggregation when DNAJA2 was knocked down, and the aggregation suppression activity of DNAJA4-CTD-II was significantly impaired (Fig. 6E). Importantly, the impairment in the aggregation suppression by DNAJA4-CTD-II upon DNAJA2 knockdown was significantly greater than the expected additive effects of DNAJA2 knockdown and DNAJA4-CTD-II overexpression alone (p=0.007 for DNAJA4-CTD-II and shDNAJA2 synergy, p=0.34 for DNAJA4-FL and shDNAJA2 synergy, see Methods), indicating that the functional effects of DNAJA4-CTD-II on TDP-43-ΔNLS–2KQ aggregation were largely dependent on DNAJA2.

Collectively, our results demonstrate that DNAJA4-CTD-II, a stress-inducible isoform of DNAJA4, forms a stress-regulated hetero-complex with DNAJA2, which functions in proteostasis regulation as a significant modulator of TDP-43 aggregation.

## Discussion

Accumulating evidence suggests that chaperone networks undergo condition-specific remodeling through changes in protein-protein interactions. However, despite the central role of chaperones in stress adaptation, surprisingly few studies have directly examined how chaperone interaction networks are reorganized under different physiological or proteotoxic conditions. Existing examples include malignant transformation ^27^, Alzheimer’s disease models ^28^, and BAG3 interactome remodeling following proteasome inhibition ^21^. Our findings further suggest that alternative isoforms contribute to chaperone networks remodeling. Previous studies showed that DNAJ isoforms can differ in their localization and aggregation-modulating activity^10,12–14,29^ Here we show that alternative isoforms can additionally remodel chaperone interaction networks through selective hetero-complex formation. Stress-dependent induction of DNAJA4-CTD-II, together with enhanced hetero-complex formation, suggests that selective incorporation of alternative isoforms provides a mechanism for dynamically reconfiguring chaperone complexes during proteotoxic stress.

Our data support a model in which DNAJA4-CTD-II functions not as an autonomous aggregation suppressor but as a regulatory component of DNAJA-containing hetero-complexes. The dependence on endogenous DNAJA2 suggests that the biologically relevant functional unit is the DNAJA2–DNAJA4-CTD-II complex rather than DNAJA4-CTD-II alone. Recent in-vitro studies have shown that DNAJA2 can directly bind the aggregation-prone prion-like domain of TDP-43 and modulate its aggregation behavior ^25,26^. As the C-terminal domains of class A DNAJ proteins have been shown to mediate substrate binding and participate directly in the recognition of aggregated model proteins ^8,30^, our findings suggest that DNAJA4-CTD-II may enhance substrate engagement, while DNAJA2-FL also provides complementary activities, likely by HSP70 recruitment to the aggregates.

Importantly, our results demonstrate that multiple distinct DNAJA isoforms differentially regulate TDP-43 aggregation. Although Class A DNAJ proteins share a highly conserved domain architecture and extensive sequence homology ^5^, our findings reveal substantial functional divergence between family members and isoforms. Notably, DNAJA1 and DNAJA2 are among the most closely related members of the Class A DNAJ family ^31^. Despite this, our study uncovered a striking functional difference between them, as DNAJA2 suppressed TDP-43 aggregation whereas DNAJA1 had no detectable modulatory effect on TDP-43 aggregates (Fig. 6B). These results reveal functional divergence among closely related class A DNAJ proteins, as well as their naturally occurring isoforms.

Together, our findings identify isoform-dependent hetero-complex assembly as a previously unexplored mechanism that expands the functional diversity of the HSP70 chaperone network. By coupling stress-regulated isoform expression with selective protein interactions, alternative DNAJ isoforms enable dynamic remodeling of proteostasis networks in response to cellular stress, and expand the functional outcomes of the network.

## Limitations of the study

The functional consequences of DNAJA4-CTD-II were examined using a cellular TDP-43 aggregation model; whether its regulation of TDP-43 extends to neuronal or disease-relevant models remains to be determined. In addition, although our network mapping identified multiple chaperone interaction modules and isoforms, functional characterization focused primarily on the DNAJA2–DNAJA4 complex. Future studies will be required to determine how isoform-dependent interaction remodeling and functional diversification extend to additional complexes identified in our network.

## Methods

### Data Sources and Genomic Annotations

Tissue-specific expression analysis was conducted using the hg38 human genome assembly, UCSC GTEx Analysis V10, and GENCODE v39 annotations. The dataset was restricted to standard human chromosomes (chr1-22, X, Y). Computational processing, statistical analysis, and data visualization were performed using R. Protein domain annotations were integrated from UniProt and Pfam, with InterPro utilized to resolve missing data. Chaperone family members were taken from Brehme et al. ^32^.

### Domain Architecture and Structural Analysis

GENCODE transcript annotations were downloaded through the UCSC Genome Browser and Ensembl, and domain annotations were obtained from the UniProt, Pfam, and InterPro databases. To determine the domain composition of each transcript, genomic coordinates of coding sequences (CDS) were intersected with domain annotations using bedtools. To perform domain assignments per isoforms, we calculated the Domain Fraction as: Domain Fraction = Intersection Length (bp)/Total Domain Length (bp). Only intersections retaining > 80% of the annotated domain length were retained. Domain nomenclature was normalized across all sources to ensure functional consistency.

Isoform Classification: Full-Length (FL) vs. Alternative (ALT) was performed as follows: For each chaperone gene (from the DNAJ, HSP70, and NEF families), a primary Full-Length (FL) isoform transcript was defined based on a four-tier selection process:

1. Functional canonicity was assigned according to the presence of the characteristic domain architecture of each chaperone family; DNAJA proteins were required to have a J-domain and zinc-finger motif, DNAJB/C proteins - a J-domain, HSP70 proteins – nucleotide-binding domain (NBD) and substrate-binding domain (SBD), and NEFs a BAG, Nucleotide_exchange, or GrpE domain. For HYOU1 and HSPH1, additional family-specific C-terminal domains were required. A “semi-canonical” classification was assigned to isoforms lacking a subset of the characteristic domains defined above, specifically in the case of HSP70 isoforms lacking either the NBD or SBD. Semi-canonical isoforms were considered candidates for FL assignment only when a chaperone gene had no isoform containing the complete canonical domain architecture.
2. Expression Threshold: Only transcripts with a minimum Transcripts Per Million (TPM) > 2 in at least one of the relevant tissues were considered as expressed and retained for further analysis.
3. Length requirement: Transcripts had to be within 85% of the maximum annotated CDS length for the respective gene.
4. Expression Dominance: Among all transcripts meeting the above criteria, the most highly expressed was designated as the FL isoform.

Alternative (ALT) isoforms included all other expressed coding transcripts. Isoforms with a CDS identical to the FL but differing in their 5’ or 3’ regions were labeled UTR_ALT. Additionally, UCSC transcripts labelled as non-coding were evaluated via ORF analysis using getorf (EMBOSS). Potential ORFs > 100 bp (starting with ATG) were retained, only if they maintained the original gene orientation, ended with a valid stop codon, and physically overlapped the CDS of the primary FL isoform. Pseudogenes DNAJB3 and HSPA7 were manually retained.

### GTEx Expression clustering and co-expression analysis

Tissue TPM values were calculated as the mean of all relevant samples. High-confidence analysis was ensured by retaining only transcripts with a minimum expression >2 TPM across the dataset, preventing low-abundance noise from biasing downstream clustering. The following tissues were excluded: Testis, Cells-Cultured fibroblasts, and Cells-EBV-transformed lymphocytes.

The expression landscape of chaperone isoforms was visualized using hierarchical clustering. The matrix was Z-score normalized per transcript across tissues. Clustering was performed using Pearson correlation as the distance metric and complete linkage. Heatmaps were annotated with metadata layers including chaperone family, isoform class (FL, ALT, UTR_ALT), and anatomical tissue groups. A secondary analysis (Fig. 2D) was restricted to chaperones validated by the LUMIER assay to cross-reference co-expression with physical interaction data.

A pairwise correlation matrix was generated for all LUMIER-validated transcripts, excluding self-interactions. We generated a global histogram of all pairwise correlations and projected the mean Pearson correlation of 24 interaction modules (Table S3), defined by their physical interaction score signatures in the LUMIER assay. To characterize the underlying co-expression distribution, we applied Gaussian Mixture Modeling (GMM), with the optimal number of components validated by the Bayesian Information Criterion (BIC).

### Plasmid and Cloning

A C-terminal FLAG-tagged library comprising 65 chaperones was generated from the ORFeome collection (GE Healthcare). The chaperone ORFs (full length (FL)) were transferred from the pDONR223 entry vectors (ORFeome) into the destination vectors pcDNA3.1-ccdb-3xFLAG-V5 or pcDNA3.1-ccdB-Nanoluc, for the bait and prey libraries respectively, using Gateway cloning.

The isoforms of DNAJA4-CTD-II (ENST00000493321.1) and DNAJA4-J-Domain (ENST00000423642.1) (Fig. 3A) and DNAJA2-CTD-I (ENST00000617000.1) (Fig. 3B) were generated from the full-length isoforms using PCR and Gateway cloning (Primers list in Supplementary Table S4). These isoforms were cloned into pAAV-CAG-ccdb-FLAG, which was generated in house.

Human TDP-43-YFP was cloned from pcDNA3.2 TDP-43 YFP (obtained through Addgene #84911^33^) into pDON221, followed by site-directed mutagenesis (Quikchange 2 Site-Directed Mutagenesis Kit, Agilent Technologies, 210518) to create two K→Q mutations at residues K145 and K192 ^22^, in addition to mutations in the nuclear localization sequence (NLS) K82A, R83A, K84A resulting in the TDP-43Δ-ΔNLS-2KQ construct, which was then cloned into pAAV-CAG-DEST backbone using Gateway cloning (primers listed in Supplementary Table S4).

For DNAJA2 knockdown, shRNA constructs against DNAJA2 or a scrambled sequence as a control (see Supplementary Table S6), were cloned into pLKO.1 - TRC cloning vector (a gift from Izhak Kehat lab), using AgeI and EcoRI sites. Oligos for DNAJA2 based on ^34^, and scrambled (in house), are listed in Supplementary Table S6. Oligos containing the AgeI/EcoRI sites underwent annealing for 45 min at RT, in annealing buffer, then diluted and ligated with the digested pLKO vector using T4 DNA ligase (NEB, M0202S), as described in https://www.addgene.org/protocols/plko/).

### Cell culture and Transfection

HEK293T and MCF7 cells were maintained in standard DMEM supplemented with 10% FBS and 1% Penicillin–Streptomycin, at 37°C in a humidified incubator with 5% CO_2_.

Transfection in HEK293T cells was done in 6-well or 10cm plates, with 2.5 µg/well or 7 µg/plate DNA respectively, 24 h after seeding, using PEI (1 mg/ml, Thermo Fisher Scientific). For the LUMIER assay, transfection was performed in 96 well plates with 200 ng/well DNA. Co-transfection was performed using a 1:1 plasmid DNA ratio.

For the generation of a stable cell line of shDNAJA2, HEK293T cells were seeded in 10 cm plates, and transfected 24 h later with 7 µg of either pLKO-shDNAJA2 or pLKO-shScrambled plasmids. Two days post transfection, selection with puromycin (Sigma-Aldrich, P8833) was applied. Knockdown of endogenous DNAJA2 was validated by Western Blotting (Fig. S5E).

For stress experiments, cells were seeded in 6-well plates and the next day were exposed to different stress conditions, as follows, for the times indicated: Thapsigargin (1 µM, Sigma-Aldrich, T9033); heat shock at 44°C; MG-132 (10 µM, Sigma-Aldrich, M7449); and sodium (meta) arsenite (200 µM, Sigma-Aldrich, S7400).

### LUMIER interactome assay

The LUMIER assay ^19^ was used to identify chaperone-cochaperone interactions, as in Taipale et al. ^19^. In brief, cells were co-transfected with bait (a FLAG tagged chaperone) and prey chaperone (a Nanoluciferase tagged chaperone), in all possible combinations for the library of 65 bait chaperones over 54 prey chaperones. After 48h, cells were harvested in LUMIER lysis buffer (50 mM HEPES pH 7.9, 0.5% Triton X-100, 5% glycerol, 150 mM NaCl, 2 mM EDTA) with protease inhibitors (a cocktail of Aprotinin, Leupeptin, Pepstatin A at 1 µg/ml each and PMSF at 400 µM). The lysate was transferred to 384-well plates (Lumitrac, Greiner Bio-One) coated with anti-FLAG M2 antibody (Sigma-Aldrich, F1804) and incubated at 4°C for 3 hr. After 7 washes with the LUMIER buffer, luminescence was measured in Infinity 200 plate reader (Tecan), using the Nano-Glo Luciferase Assay reagent (Promega, N1120). After additional washes, HRP-conjugated anti-FLAG antibody (Abcam, ab1238) was added in ELISA buffer (PBS, 2% goat serum, 5% Tween 20). Plates were incubated for 1.5 h, washed in PBS+0.05% Tween 20 with a plate washer, and ELISA signal was detected with 3,3’,5,5’-tetramethylbenzidine (TMB) substrate.

Each LUMIER experiment was performed with n=2 replicates, and Interaction Scores (Prey/Bait, using measured luciferase activity / FLAG ELISA levels) were averaged to generate an IS vector per prey per experiment. For each prey chaperone, this was repeated in multiple biologically independent experiments, with N=3-5 for most chaperone preys (except DNAJB12, HSPA14 and HSPA2 which had 9/6/6 replicates). Each individual prey IS vector was z-score normalized, and all IS for all independent replicates were then averaged to generate the final IS, presented in Fig. 2A (see Table S7). The final interactome network contained 54 prey chaperones and 65 bait chaperones. In all cases full-length isoforms were used; In the case of five chaperones: HSPH1, DNAJB2, HSPA9, DNAJA3, and DNAJC6, FL isoforms selected for tissue expression analysis differed from the ORFeome isoforms, as those had higher mean expression levels across tissues. Nonetheless, in all five cases, they were canonical and contained all functional protein domains, and differed in either their very N- or C-termini. With the exception of HSPA9, all tissue expression-defined FL isoforms clustered in the same tissue expression cluster as the ORFeome FL isoform.

Co-transfections of bait-prey pairs were performed for all prey proteins, except in the case of Hspa8 and Hspa1a, for which stable cell lines expressing the luciferase-tagged preys were used. Interaction scores were clustered using hierarchical clustering, z-score normalized IS between -1 and 1 shown as white in Fig. 2A. The complete IS score data is provided in Supplementary Table S7.

### Co-immunoprecipitation

Co-immunoprecipitation (co-IP) was performed in LUMIER buffer using Anti-FLAG M2 magnetic beads (Sigma-Aldrich, M8823). Cells were transfected with plasmids of interest, and harvested 48h post-transfection, in LUMIER buffer with proteasome inhibitors, on ice. Lysates were centrifuged at 12,000g for 10 min at 4°C, and the clarified supernatants were transferred to fresh tubes for co-IP and incubated with beads overnight at 4°C. After four washes with LUMIER buffer, bound proteins were eluted with 50 mM glycine (pH 2.5). Equal volumes of eluates were resolved on a 12% SDS-PAGE gel and immunoblotted using anti-FLAG (Sigma-Aldrich, F1804) or antibodies for the co-precipitated proteins, including: anti-GFP (4B10) (MBL #2955), anti-HSC70/HSP70 (Enzo, ADI-SPA-820-F), anti-HSPA6 (Santa Cruz, sc-374589) or anti DNAJA2 (Santa Cruz, sc-136515), as shown in (Fig. 2, 3, 6). For sodium arsenite experiment (Fig. 5A,B), cells were treated with 200 µM sodium arsenite (Sigma-Aldrich, S7400) for 2h, after which the medium was replaced and cells were allowed to recover for an additional 2h, before co-IP.

### PulSA method

To quantify the aggregation of TDP-43 by flow cytometry we applied the PulSA method ^23^, as previously described ^10^. HEK293T cells were transfected with a total amount of 2.5 ug DNA per well, in a 6 well plate, including TDP-43Δ-ΔNLS-2KQ and potential modifier chaperones, or DsRed as control. After 48h cells were collected in growth media, in the presence of DAPI (Sigma-Aldrich, D9542) to evaluate cell viability, and subjected to FACS analysis. Cells with aggregates were separated from those with diffused fluorescence by plotting GFP peak height (GFP-H) versus peak width (GFP-W). Gates define GFP negative cells (GFP^-^), aggregate-containing cells (AGG^+^), or cells with diffused GFP fluorescence (AGG^-^). (Fig. S5B). The Aggregation Modulation Score was calculated as log_2_(%AGG^+^ (modifier) / %AGG^+^ (Control)). Confidence intervals (95% CI, appear as dashed lines in Fig. 6B, D) were calculated according to the variations between different control (DsRed) replicates through different experiments, such that they represent twice the standard deviation of Aggregation Modulation Scores calculated for the population of DsRed. Chaperones were called as significant modulators if their mean Aggregation modulation score +/− SEM was below/above the 95% CI in the case of a negative/positive score respectively. An empirical *p*-value was indicated as *p* < 0.05 (*) if the score value was below/above the 95% CI (i.e. twice the STD), and *p* < 0.003 (***) if the score was below the 99.7% CI (i.e., 3 times STD). TDP-43 CI = +/-0.172 was determined using a total of n=30 replicates for Fig. 6B, and CI = +/-0.166 was determined using a total n=22 for Fig. 6E in the pLKO-shScramble cell lines.

### qPCR

RNA was extracted using the Quick-RNA Miniprep Kit (Zymo, R1055), which included on-column DNase treatment for 15 min. We additionally followed with a second DNase treatment using RQ1 RNase-Free DNase (Promega, M6101), according to the manufacturer’s protocol. cDNA was synthesized using LunaScript® RT SuperMix (NEB, M3010L). qPCR was performed using Luna® Universal qPCR Master Mix (NEB, M3003E) with primers specific for the DNAJA4 and DNAJA2 isoforms and HPRT (primer sequences are listed in Supplementary Table S5). Genomic DNA, extracted from MCF7 cells using Quick-DNA Miniprep Kit (Zymo, D3024) was used for standardization in qPCR experiments as in Shalgi et. al., 2014 ^35^. qPCR products were calculated as absolute DNA copy numbers relative to the standard curve of genomic DNA ranging from 50ng (14,300 copies) to 0.4ng (114 copies) ^36^. No-RT (no reverse transcriptase) control reactions were included in all experiments, and amplification from residual genomic DNA was excluded.

### Immunofluorescence staining

Cells were cultured on coverslips, in 6 well plates and transfected as described above. Two days post transfection, cells were fixed in 4% paraformaldehyde in PBS, permeabilized with 0.5% Triton-x100 in IF buffer (5% FCS, 2% BSA in PBS), then immunostained with the anti-FLAG antibody (Sigma-Aldrich, F1804), followed by the secondary antibody AlexaFluor 647 Donkey Anti-Mouse (Jackson ImmunoResearch Labs. 715-605-151), and DAPI staining of nuclei. Images were acquired using laser scanning confocal microscope (LSM900).

### Statistical analysis of synergy

Synergy between DNAJA4-FL/DNAJA4-CTD-II overexpression and DNAJA2 knockdown was evaluated separately for each overexpression construct using a two-way ANOVA. The model included the main effects of overexpression (control vs. overexpression), knockdown (control vs. knockdown), and their interaction. The interaction term was used as the statistical test for synergy, with a significant interaction indicating that the combined effect of overexpression and knockdown differed from the additive effects of the individual perturbations. Statistical significance was assessed using the p-value of the interaction term. The interaction term was calculated as: mean(OX+KD) − mean(OX) − mean(KD) + mean(Control), while for DNAJA4-FL interaction term was 0.1 and p=0.34 and for DNAJA4-CTD-II the interaction term was 0.32 and p=0.007, indicating significant deviation from the expected score assuming independence.

## Acknowledgements

This work was supported by the Israel Science Foundation (ISF, 694/18 and 1494/24). We thank the Prince Center for Neurodegenerative disorders of the brain, and the Rappaport Family Institute for Research in the Medical Sciences for their funding support. NA was supported by the Israeli VATAT national doctoral fellowship.

## Author contributions

The project was conceived by RS. RS, TK, and FLA designed all experiments. TK, ABN and FLA performed all experiments. NA and AM performed all data analysis. RS and TK wrote the manuscript.

## Data Availability

The datasets generated during the current study are included in this published article and its supplementary information files. Raw data used to generate Aggregation Modulation Scores are available from the corresponding author upon request.

## Declaration of generative AI and AI-assisted technologies in the manuscript preparation process

During the preparation of this work, the authors used ChatGPT for grammar and language editing checks. The authors reviewed and edited the output as needed and take full responsibility for the content of the published article.

